# Computational Analysis of Antioxidant Effects on Stress Response Mechanisms in *Candida albicans*

**DOI:** 10.64898/2026.10.31.762764

**Authors:** K R Harshitha, G Sriprada, S Atiya Naaz, Shivakumar, Rashmi Shenoy, Savithri Bhat

## Abstract

Candida albicans was used as a eukaryotic model to study aging and oxidative stress response pathways focused on two NAD□ dependent deacetylases, Sir2 and SIRT3. Gene expression analysis under sublethal hydrogen peroxide treatment showed significant upregulation of both Sir2 and SIRT3, connecting them to stress adaptation. Natural ligands known for antioxidant properties were docked against Sir2 and SIRT3 structures. Binding free energy of most components fall in the range of −5.0 to −11 kcal/mol. The best candidates pterostilbene, quercetin and naringenin formed stable hydrophobic interactions and hydrogen bonds within catalytic and NAD□ binding pockets confirmed by visualization. Detailed ADME profiling confirmed that most ligands passed Lipinski’s rule of five and indicated great gastrointestinal absorption. PASS (Prediction of Activity Spectra for Substances) and SOM (Sites of Metabolism) predicts antioxidants, anti-aging properties, toxicity profiles, with key metabolic hotspots. This integrated *in-silico* approach highlights naringenin and quercetin as promising modulators of Sir2/SIRT3 mediated metabolic pathways in Candida albicans, warranting further validation with the help of *in-vitro* and *in-vivo* methods.

## I. INTRODUCTION

AGING is a natural biological process characterized by a gradual decline in cellular function, increased susceptibility to environmental and physiological stressors, and reduced ability to maintain homeostasis. Oxidative stress is one of the primary causes of cellular aging, since it leads to the accumulation of reactive oxygen species (ROS), which induce damage to critical biomolecules such as DNA, proteins, and lipids. This damage leads to impaired cellular functions, senescence, and ultimately, cell death, thus promoting aging and related pathologies *(Mladenov et al., 2023)*. In this context, understanding the molecular pathways that govern oxidative stress responses is essential to develop strategies to mitigate aging and its associated disorders.

*Candida albicans*, a dimorphic fungal pathogen, has emerged as an important model organism for investigating oxidative stress and aging-related processes owing to its conserved stress response pathways and ability to survive in host environments *(Seneviratne et al., 2008; Tscherner et al., 2015)*. Notably, antioxidant defenses in *C. albicans*, including enzymatic systems and metabolic adaptations, play a major role in its survival during oxidative insults, especially in its pathogenesis and biofilm formation *(González-Párraga et al., 2003)*. These antioxidant responses are closely related to cellular longevity and stress tolerance mechanisms that are similar to higher eukaryotes.

Of particular interest are the sirtuin family proteins, such as SIRT3 and Sir2, NAD□-dependent histone deacetylases that coordinate cellular responses to oxidative stress and control longevity by means of metabolic regulation, DNA repair, and chromatin remodeling. These sirtuins modulate gene expression patterns and enhance cellular resistance to oxidative damage, thus representing promising molecular targets for anti-aging interventions *(Ogura et al., 2021)*. Computational analysis, consisting of gene characterization, molecular docking of sirtuin receptors with various antioxidant ligands, and *in-silico* pharmacokinetics and toxicity predictions (ADME, PASS, SOM), helps identify the therapeutic significance of antioxidants in influencing sirtuin-mediated pathways.

This project employs a problem-based and project-based learning framework to explore the impact of antioxidants on stress response pathways in C. albicans. By combining gene analysis of SIRT3 and Sir2 with receptor-ligand docking of 25 antioxidants compounds and subsequent *in-silico* ADME and activity spectrum predictions, we aim to describe how oxidative stress-induced aging can be mitigated at a molecular level. In addition to improving our knowledge on fungal stress resistance and aging, this research also helps in identifying potential antioxidant interventions that may be used in broader anti-aging applications.

## II. Tools, Dataset and Methodology

### A. Ligand selection and preparation

The chemical structures of phytochemicals or drug-like compounds were obtained from the PubChem database (https://pubchem.ncbi.nlm.nih.gov/) in SDF format. These structures were then converted into the appropriate file format (e.g., PDB or PDBQT) using Open Babel for further analysis

### B. ADME analysis and drug-likeliness prediction

Absorption, distribution, metabolism, excretion analysis is crucial to understand the pharmacokinetics of any ligand inside system. Parameters such as Lipinski’s Rule of Five, gastrointestinal (GI) absorption, bioavailability score, and blood-brain barrier (BBB) permeability were assessed (*Morcoss et al., 2020*). Ligands considered were curcumin, procyanidin, ascorbic acid, luteolin, ellagic acid, fisetin, EGCG, pterostilbene, vitamin E, resveratrol, genistein, flavonoids, naringenin, beta-carotene, and lipoic acid which are bioactive compounds found in various fruits, vegetables and other food materials. For natural compounds like the ones considered here, ADME analysis helps to identify potential bioavailable soluble and effective drug candidates.

The SMILES file format of all the ligands considered was obtained using PubChem (PubChem). To evaluate the pharmacokinetic properties and drug-likeness of the selected compounds, SwissADME (http://www.swissadme.ch/) was used. The results obtained were analysed (*Opo et al., 2021*).

### C. Functional gene analysis of SIRT3 and Sir2

To validate the selection of the receptors SIRT3 (from *Homo sapiens*) and Sir2 (from *Candida albicans*), interaction network and pathway enrichment analysis were performed using STRING (https://string-db.org/) and ShinyGO (https://bioinformatics.sdstate.edu/go/) (*Steven et al., 2021*)

### D. Selection and preparation of receptors

Two target proteins were selected based on their functional gene analysis their relevance in ageing and oxidative stress (*Seneviratne et al., 2008*).

### E. Molecular docking

Molecular docking studies were performed using AutoDock, a widely used tool for predicting ligand binding affinity and interaction. The prepared ligand and protein files were used as input, and a grid box was defined around the active site of the protein. The exhaustiveness was set to 8 or higher to improve accuracy (*Kechi et al., 2024*).

The molecular docking study assessed natural functional food compounds for their potential anti-aging and anti-stress effects through interactions with key proteins involved in these processes, namely the human SIRT3 receptor and the SIRT1 (Sir2) receptor. Ligands such as curcumin, procyanidin, ascorbic acid, luteolin, ellagic acid, fisetin, EGCG, pterostilbene, vitamin E, resveratrol, genistein, flavonoids, naringenin, beta-carotene, and lipoic acid are bioactive compounds found in various fruits, vegetables, and spices. These compounds are known for their antioxidant and anti-inflammatory properties, which help mitigate oxidative stress and inflammation, major contributors to aging and stress-related damage. Their binding to SIRT3 suggests modulation of immune and inflammatory pathways, while interaction with Sir2 — a crucial regulator of cellular longevity and stress resistance — indicates their potential to promote healthy aging and enhance cellular resilience.

### F. Visualization of ligand and receptor interaction

Following molecular docking, the resulting ligand–receptor complexes were visualized to examine the molecular interactions between the ligands and the receptor binding sites. Visualization tool PyMOL was employed to analyse the binding modes and identify key amino acid residues involved in ligand recognition and stabilization (*Opo et al., 2021*).

The docked complexes were loaded into the visualization software, and the interactions were assessed based on proximity (typically within 4 Å) and type of interaction, including hydrogen bonding, hydrophobic contacts, electrostatic interactions, and π–π stacking. For each complex, the amino acid residues forming direct contacts with the ligand were identified and recorded. The visualization helped in understanding the orientation and fit of the ligands within the binding pocket, as well as the role of specific residues in ligand accommodation. This analysis provided structural insights into receptor–ligand complementarity, which is useful for evaluating binding specificity and for guiding future *in vitro* or *in-silico* studies.

### G. PASS, SOM prediction and chemical interaction analysis

The canonical SMILES from the PubChem server, https://pubchem.ncbi.nlm.nih.gov/, were utilized by the PASS-Way2Drug server, http://www.pharmaexpert.ru/passonline/, to assess an organic drug-like molecule’s potential for biological functions. A web-service for *in-silico* SOMP, PASS (Prediction of Activity Spectra for Substances) (http://www.way2drug.com/PASSonline) and LMNA descriptors are the foundations for prediction (*Waykar & Kumarapillai, 2024*). Biovia Discovery Studio Visualizer v21.1.0.20298 was used to view the molecular interactions in both 2D and 3D.

## III. Results and discussion

### A. Gene analysis

To strengthen the rationale behind selecting SIRT3 and Sir2 as receptors, a functional enrichment analysis was performed using STRING (https://string-db.org/) and ShinyGO (http://bioinformatics.sdstate.edu/go/).

#### 1) STRING analysis

**Fig 1.**
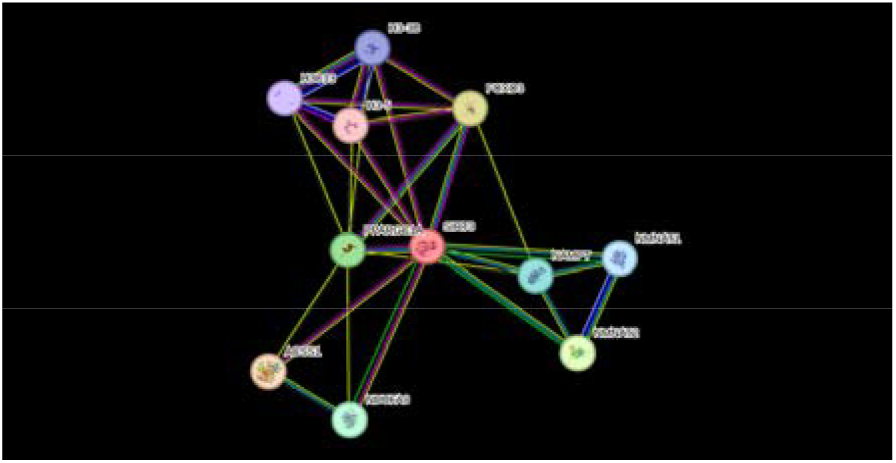
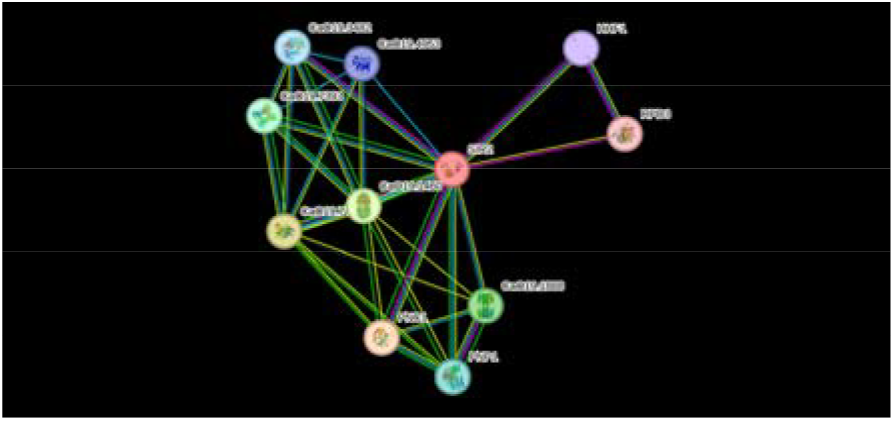
Predicted functional partners of SIRT3 and Sir2.

**TABLE I.** PREDICTED FUNCTIONAL PARTNERS OF SIRT3 (STRING ANALYSIS)

| **Gene** | **Description** | **Score** |
|---|---|---|
| **ACSS1** | Acetyl-CoA synthetase; mitochondrial energy metabolism | 0.994 |
| **FOXO3** | Transcription factor; binds DNA sequences | 0.987 |
| **NMNAT2** | NAD□ synthesis enzyme; neuronal NAD□ metabolism | 0.983 |
| **PPARGC1A** | Coactivator of mitochondrial and oxidative genes | 0.978 |
| **NDUFA9** | Mitochondrial complex I subunit | 0.976 |
| **NAMPT** | NAD□ biosynthesis; rate-limiting enzyme | 0.970 |
| **NMNAT1** | NAD□ synthesis enzyme; nuclear | 0.950 |
| **H3-3B** | Histone variant H3.3; nucleosome regulation | 0.950 |
| **H3C13** | Histone H3.2; chromatin compaction | 0.950 |
| **H3-5** | Histone H3.3C; nucleosome structure | 0.950 |

**TABLE II.** PREDICTED FUNCTIONAL PARTNERS OF SIR2 (STRING ANALYSIS)

| **Gene/ID** | **Short Description** | **Score** |
|---|---|---|
| **PNC1** | Nicotinamidase | 0.935 |
| **CaO19.7499** | NMN adenylyltransferase family | 0.932 |
| **CaO19.1460** | NAD (+) synthetase family | 0.920 |
| **CaO19.1888** | Uridine nucleosidase | 0.915 |
| **CaO19.7393** | NADH/NAD (+) kinase | 0.915 |
| **PNP1** | Purine nucleoside phosphorylase | 0.912 |
| **CaO19.3482** | NAD (+) diphosphatase | 0.904 |
| **CaO19.4953** | NMN adenylyl transferase | 0.900 |
| **HHF1** | Histone H4; nucleosome structure | 0.887 |
| **RPD3** | Histone deacetylase (HDAC family) | 0.882 |

#### 1) ShinyGo gene analysis

Enrichment analysis of SIRT3 and Sir2 and their predicted functional partners revealed significant overrepresentation in metabolic and aging-related pathways:

**TABLE III.** SIGNIFICANTLY ENRICHED PATHWAYS AMONGST THE GENES INTERCATING WITH SIRT3 AND SIR2.

| Pathway | Fold Enrichment | Relevance |
|---|---|---|
| Nicotinate and nicotinamide metabolism | 231.1 | NAD□ biosynthesis |
| Longevity regulating pathway | 6.7 | Aging and lifespan regulation |
| AMPK signaling pathway | 0.4 | Energy homeostasis |
| Metabolic pathways | 0.1 | General cellular metabolism |
| Nicotinate and nicotinamide metabolism | 47.9 | NAD□ biosynthesis |
| Longevity regulating pathway- multiple species | 00.2 | Aging and lifespan regulation |

**Fig 2.**
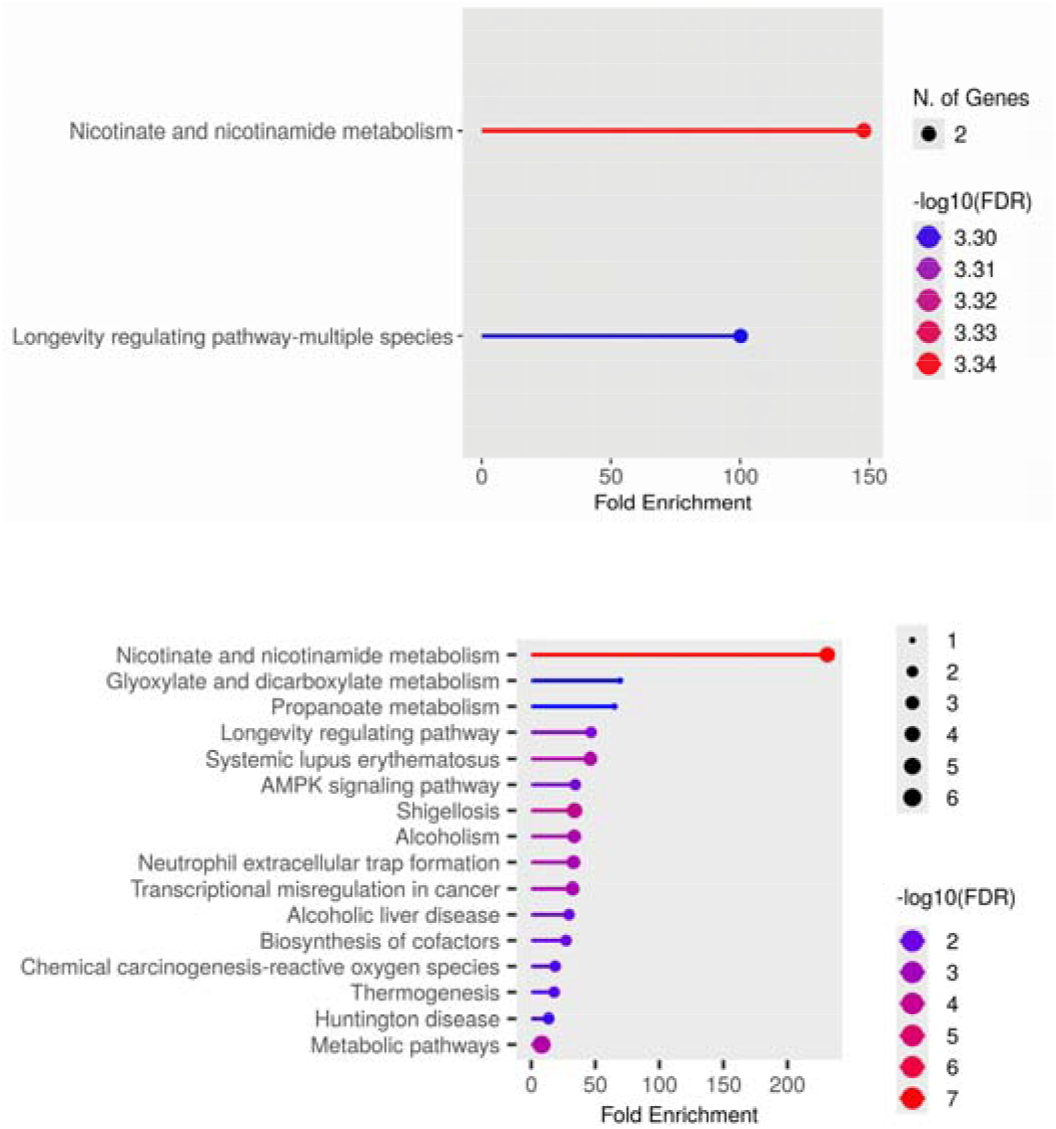
Pathway enrichment plots of Sir2 and SIRT3

These results support the role of SIRT3 in energy metabolism, stress response, and lifespan regulation, in agreement with previous findings Sir2-associated genes are functionally focused, primarily enriched in NAD□ metabolism and longevity pathways, which aligns with the known biological role of Sir2 in aging and energy regulation.

For SIRT3 receptor, comparison with the human genome showed that most SIRT3-associated genes are protein-coding. Gene features such as number of exons, transcript isoforms, UTR lengths, and GC content showed no significant deviation from genomic averages (Chi-square P > 0.6).

This supports the genomic stability of SIRT3 targets and validates their use in downstream analyses.

Unlike broad genomic variability, Sir2 targets converge on conserved biological processes, with no evidence of widespread pathway divergence. The limited number of significantly enriched pathways (only 2) suggests that Sir2-associated genes maintain tight functional clustering, reducing noise and increasing biological specificity.

**Fig 3.**
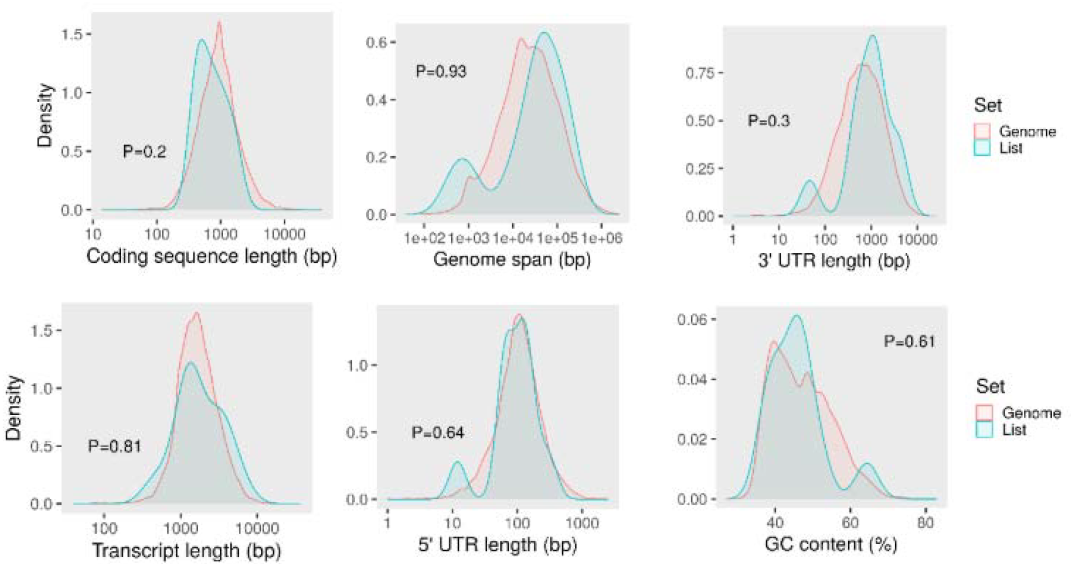
Distribution of Gene Characteristics in SIRT3-Associated Genes vs Genome

Gene enrichment analysis revealed that both SIRT3 (human) and Sir2 (Candida albicans) are functionally linked to key aging-related pathways, including nicotinate and nicotinamide metabolism and the longevity regulating pathway. These pathways are essential for maintaining NAD□ balance, mitochondrial health, and stress resistance — all central features of cellular aging. The presence of conserved interactions with chromatin-associated proteins and NAD□-dependent enzymes in both species supports the idea that SIRT3 and Sir2 share a conserved role in promoting longevity and metabolic regulation. This validates their selection as cross-species targets for studying anti-aging mechanisms and mitochondrial function.

### B. Molecular docking

Molecular docking studies were conducted using SIRT3(Sirtuin 3) from human & Sir2 from *C*.*albicans* to evaluate the binding affinity of various natural compounds. The binding energy values (in kcal/mol) are summarized in the table below:

**TABLE IV.** DOCKING SCORES OF LIGANDS WITH HUMAN SIRT3 (5D7N) & *C. albicans* SIR2(2HJH) RECEPTOR.

| **S. No.** | **Ligand** | **SIR2 (C. albicans)** | **SIRT3 (Human)** |
|---|---|---|---|
| **1** | Curcumin | -5.57 | -14.17 |
| **2** | Resveratrol | -8.30 | -10.49 |
| **3** | Flavonoid | -4.71 | -9.75 |
| **4** | EGCG | -5.46 | -10.78 |
| **5** | Fisetin | -6.85 | -9.99 |
| **6** | Genistein | -4.67 | -9.35 |
| **7** | Ellagic acid | -5.45 | -9.85 |
| **8** | Naringenin | -8.03 | -10.5 |
| **9** | Luteolin | -6.94 | -10.94 |
| **10** | Pterostilbene | -5.74 | -10.05 |
| **11** | Spermidine | -1.27 | -3.97 |
| **12** | Beta Carotene | -3.93 | -12.61 |
| **13** | Coenzyme Q10 | +0.05 | +0.09 |
| **14** | Caffeic acid | -4.05 | -6.68 |
| **15** | Alpha-tocopherol | -4.01 | -11.7 |
| **16** | Ascorbate 2 Phosphate | -4.56 | -8.53 |
| **17** | N-acetylcysteine | -3.59 | -4.92 |
| **18** | Trolox | -4.68 | -9.37 |
| **19** | Lipoic acid | -5.27 | -6.4 |
| **20** | Vitamin E | -7.12 | -12.54 |
| **21** | Sulphoraphane | -3.53 | -5.56 |
| **22** | Procyanidin B2 | -9.36 | -16.57 |
| **23** | Epigallocatechin gallate | -5.54 | -6.99 |
| **24** | Ascorbic acid | -8.26 | -9.23 |

### C. Visualization

**Fig 4.**
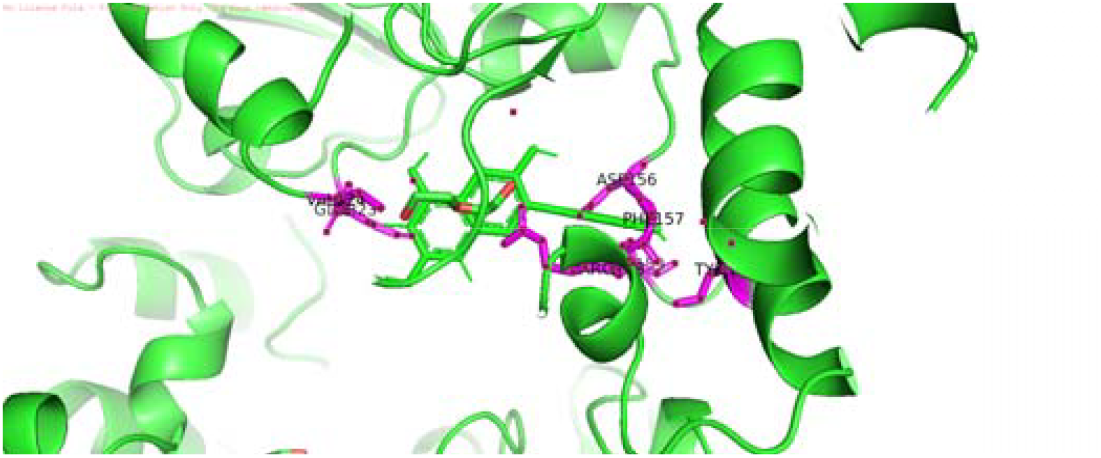
Visualization of SIRT3-Quercetin complex

Amino Acid Residues Involved in Ligand Binding to SIRT3 Receptor: ALA146, ASP156, PHE157, ARG158, SER159, SER162, TYR165, SER321, GLU323, VAL324

**Fig 5.**
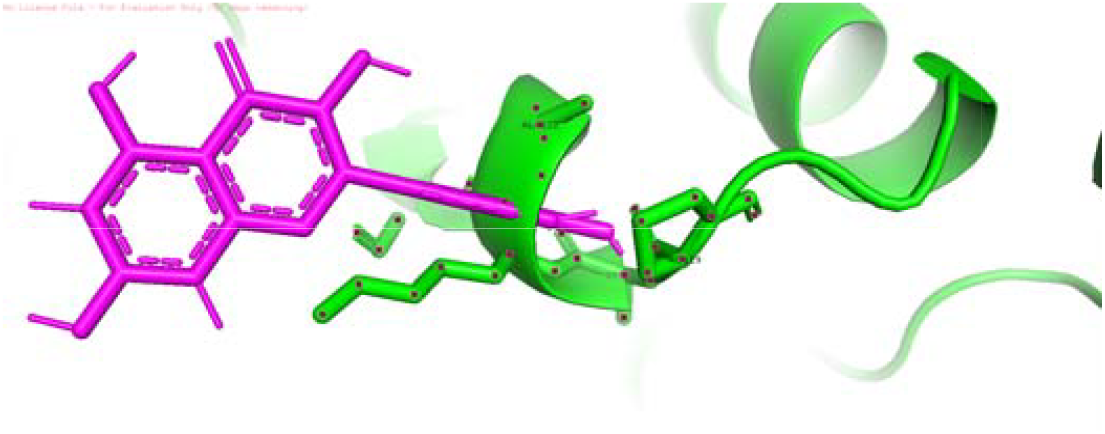
Visualization of Sir2-Quercetin complex

Amino Acid Residues Involved in Ligand Binding to SIRT3 Receptor: TYR118, PRO119, LYS120, LYS121, ALA122, VAL123, GLU124

**Fig 6.**
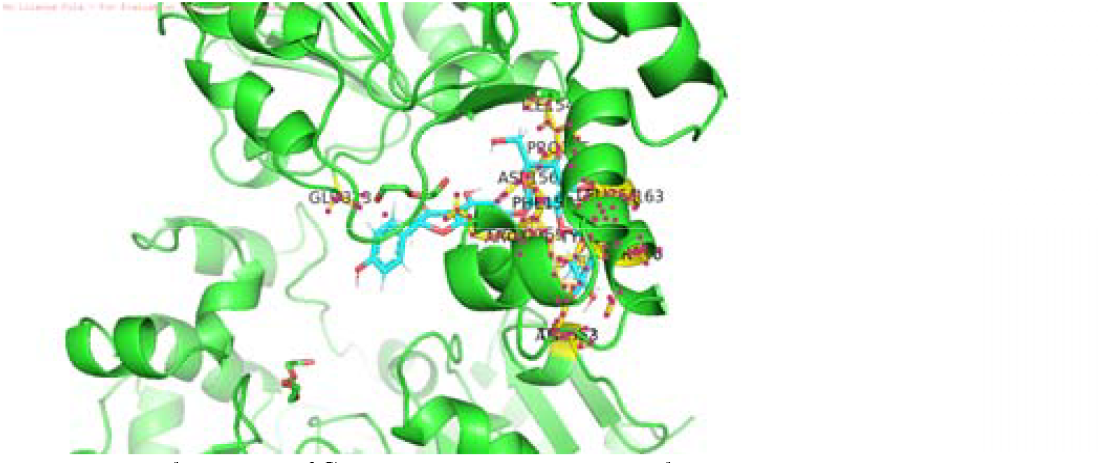
Visualization of SIRT3-Naringenin complex

Amino Acid Residues Involved in Ligand Binding to SIRT3 Receptor: VAL141, VAL142, MET 43, PHE309, LEU315, PHE327, ALA328, SER329, LEU330, THR331, GLU332, ALA333, VAL334, ARG335, SER336, ARG340, ARG356

**Fig 7.**
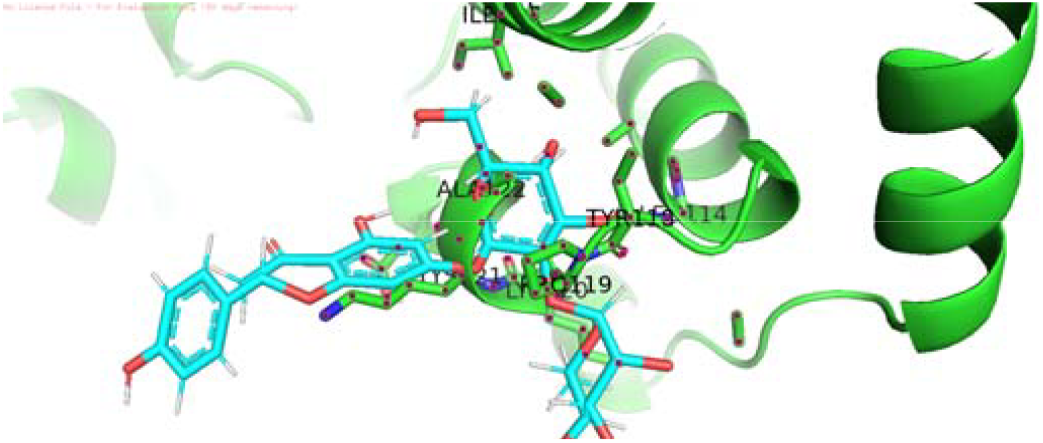
Visualization of Sir2-Naringenin complex

Amino Acid Residues Involved in Ligand Binding to SIR2 Receptor: THR245, GLY246, ALA247, GLY248, ILE249, ARG259, GLN328, ASN329, HIS348, ILE429, GLY430, THR431, SER432, LEU433, LYS434, VAL435, GLU436

### D. ADME analysis

**TABLE V.** ADME RESULTS OF LIGANDS UNDER CONSIDERATION.

| Molecule | ESOL Class | Ali<br>Log S | GI<br>absorption | BBB<br>permeant | log Kp<br>(cm/s) |
|---|---|---|---|---|---|
| Curcumin | Soluble | -4.83 | High | No | -6.28 |
| Resveratrol | Soluble | -4.07 | High | Yes | -5.47 |
| Flavanone | Moderately soluble | -4.14 | High | Yes | -5.34 |
| EGCG | Soluble | -4.91 | Low | No | -8.27 |
| Fisetin | Soluble | -3.93 | High | No | -6.65 |
| Genistein | Soluble | -4.23 | High | No | -6.05 |
| Ellagic acid | Soluble | -3.66 | High | No | -7.36 |
| Naringenin | Soluble | -3.99 | High | No | -6.17 |
| Luteolin | Soluble | -4.51 | High | No | -6.25 |
| Pterostilbene | Moderately soluble | -4.29 | High | Yes | -5.18 |
| Spermidine | Highly soluble | 0.15 | High | No | -7.9 |
| Beta carotene | Highly soluble | 0.15 | High | No | -7.9 |
| Coenzyme q10 | Highly soluble | 0.15 | High | No | -7.9 |
| Caffeic acid | Very soluble | -2.38 | High | No | -6.58 |
| Alpha tocopherol | Poorly soluble | -11.27 | Low | No | -1.33 |
| Ascorbate-2-phosphate | Highly soluble | 0.08 | Low | No | -9.96 |
| N-acetyl-cystein | Very soluble | -2.13 | High | No | -7.04 |
| Trolox | Soluble | -3.9 | High | Yes | -5.81 |
| Lipoic acid | Very soluble | -3.14 | High | No | -6.37 |
| Vitamin E | Poorly soluble | -11.27 | Low | No | -1.33 |
| Sulforaphane | Very soluble | -2.78 | High | No | -6.32 |
| Procyanidin B2 | Moderately soluble | -6.65 | Low | No | -8.15 |
| Epigallocatechin | Soluble | -2.29 | High | No | -8.17 |
| Quercetin | Soluble | -3.91 | High | No | -7.05 |
| Ascorbic acid | Highly soluble | -0.1 | High | No | -8.54 |

**TABLE VI.** OTHER IMPORTANT ASPECTS FROM ADME ANALYSIS.

| Molecule | Lipinski violation | Bioavailability Score | Lead Likeness violation | Consensus Log P |
|---|---|---|---|---|
| Curcumin | 0 | 0.55 | 2 | 3.03 |
| Resveratrol | 0 | 0.55 | 1 | 2.48 |
| Flavanone | 0 | 0.55 | 1 | 2.84 |
| EGCG | 2 | 0.17 | 1 | 0.95 |
| Fisetin | 0 | 0.55 | 0 | 1.55 |
| Genistein | 0 | 0.55 | 0 | 2.04 |
| Ellagic acid | 0 | 0.55 | 0 | 1 |
| Naringenin | 0 | 0.55 | 0 | 1.84 |
| Luteolin | 0 | 0.55 | 0 | 1.73 |
| Pterostilbene | 0 | 0.55 | 1 | 3.31 |
| Spermidine | 0 | 0.55 | 1 | 0.12 |
| Beta carotene | 0 | 0.55 | 1 | 0.12 |
| Coenzyme q10 | 0 | 0.55 | 1 | 0.12 |
| Caffeic acid | 0 | 0.56 | 1 | 0.93 |
| Alpha tocopherol | 1 | 0.55 | 3 | 8.29 |
| Ascorbate-2-phosphate | 0 | 0.11 | 0 | -2.3 |
| N-acetyl-cystein | 0 | 0.56 | 1 | -0.07 |
| Trolox | 0 | 0.85 | 0 | 2.47 |
| Lipoic acid | 0 | 0.56 | 1 | 2.04 |
| Vitamin E | 1 | 0.55 | 3 | 8.27 |
| Sulforaphane | 0 | 0.55 | 1 | 1.92 |
| Procyanidin B2 | 3 | 0.17 | 1 | 1.39 |
| Epigallocatechin | 1 | 0.55 | 0 | 0.42 |
| Ascorbic acid | 0 | 0.56 | 1 | -1.42 |
| Quercetin | 0 | 0.55 | 0 | 1.23 |

### E. PASS, SOM prediction and chemical interaction analysis

Compounds with a low Pi (<0.5) and a high Pa (>0.7) for a desired activity are regarded as possible leads in PASS analysis. The Pa and Pi values help quantify the predictions into probability scores. All 25 ligands had low probabilities of inactivity (Pi < 0.005) and high probabilities of activity (Pa > 0.7), according to PASS prediction analysis, indicating their potential as bioactive candidates. These were found to be membrane integrity agonists and inhibitors of NADPH peroxidase, HIF1A expression, HMOX1 expression and Chlordecone reductase, indicating potential roles in detoxification and oxidative stress protection. Among them, the most favorable profiles were those of Luteolin (Pa: 0.978, Pi: 0.001), Caffeic acid (Pa: 0.977, Pi: 0.001), Lipoic acid (Pa: 0.977, Pi: 0.001), and Naringenin (Pa: 0.918, Pi: 0.004).

SOM prediction helps understanding the substitution of atoms with another for higher binding affinity through the Delta P values. The ligands tested showed no negative values in the SOM prediction results, indicating that atoms can be substituted and experimented and tested for higher affinity. Ellagic acid (ΔP = 70.1), fisetin (ΔP = 67.5), luteolin (ΔP = 66.7), and quercetin (ΔP = 66.7), among other compounds, were further highlighted by SOM prediction results as structurally modifiable for increased binding affinity.

With strong biological activity and the potential for modification, luteolin, fisetin, quercetin, ellagic acid, and naringenin were chosen as promising antioxidant leads for additional experimental validation based on the combined PASS and SOMP analysis

**TABLE VII.** PASS PREDICTION OF DRUG MOLECULES UNDER STUDY.

| SL No | Ligand | Pa | Pi | Predicted Activity |
|---|---|---|---|---|
| **1** | Curcumin | 0.936 | 0.003 | Feruloyl esterase inhibitor |
| **2** | Resveratrol | 0.941 | 0.004 | Membrane integrity agonist |
| **3** | Flavanone | 0.962 | 0.003 | Membrane integrity agonist |
| **4** | EGCG | 0.969 | 0.002 | HMOX1 expression enhancer |
| **5** | Fisetin | 0.943 | 0.001 | Antimutagenic |
| **6** | Genistein | 0.939 | 0.004 | HIF1A expression inhibitor |
| **7** | Ellagic acid | 0.907 | 0.005 | Ubiquinol-cytochrome-c reductase inhibitor |
| **8** | Naringenin | 0.918 | 0.004 | Chlordecone reductase inhibitor |
| **9** | Luteolin | 0.978 | 0.001 | Chlordecone reductase inhibitor |
| **10** | Pterostilbene | 0.935 | 0.005 | Membrane integrity agonist |
| **11** | Spermidine | 0.941 | 0.004 | Membrane integrity agonist |
| **12** | Beta carotene | 0.985 | 0.000 | Phosphatidylcholine-retinol O-acyltransferase inhibitor |
| **13** | Coenzyme Q10 | 0.954 | 0.002 | Reductant |
| **14** | Caffeic acid | 0.977 | 0.001 | Feruloyl esterase inhibitor |
| **15** | Alpha Tocopherol | 0.970 | 0.002 | Lipid peroxidase inhibitor |
| **16** | Ascorbate-2-phosphate | 0.962 | 0.001 | NADPH peroxidase inhibitor |
| **17** | N-acetylcysteine | 0.945 | 0.002 | NADPH peroxidase inhibitor |
| **18** | Trolox | 0.964 | 0.003 | CYP2C12 substrate |
| **19** | Lipoic acid | 0.977 | 0.001 | (S)-2-hydroxy-acid oxidase inhibitor |
| **20** | Vitamin E | 0.970 | 0.002 | Lipid peroxidase inhibitor |
| **21** | Sulphoraphane | 0.948 | 0.002 | CYP2E1 inhibitor |
| **22** | Procyanidine B2 | 0.968 | 0.002 | Membrane integrity agonist |
| **23** | Epigallocatechin | 0.975 | 0.001 | HMOX1 expression enhancer |
| **24** | Ascorbic acid | 0.928 | 0.003 | Antioxidant |
| **25** | Quercetin | 0.962 | 0.001 | Peroxidase inhibitor |

**TABLE VIII.** SOM PREDICTION OF DRUG MOLECULES UNDER STUDY.

| **Ligand** | **Atom Number** | **Rank** | **ΔP** |
|---|---|---|---|
| Curcumin | C.14 | 1 | 43.4 |
| Resveratrol | C.9 | 1 | 60.0 |
| EGCG | C.21 | 1 | 57.0 |
| Genistein | C.11 | 1 | 66.6 |
| Fisetin | C.15 | 1 | 67.5 |
| Ellagic acid | C.1 | 1 | 70.1 |
| Naringenin | C.6 | 1 | 42.3 |
| Luteolin | C.15 | 1 | 66.7 |
| Pterostilbene | C.1 | 1 | 51.5 |
| Spermidine | C.7 | 1 | 32.8 |
| Beta-carotene | C.14 | 1 | 50.4 |
| Coenzyme Q10 | C.14 | 1 | 40.9 |
| Caffeic acid | C.1 | 1 | 68.0 |
| Alpha-tocopherol | O.29 | 1 | 7.0 |
| Ascorbate 2-phosphate | C.1 | 1 | 54.0 |
| N-acetylcysteine | C.1 | 1 | 50.3 |
| Trolox | C.1 | 1 | 57.3 |
| Vitamin E | S.3 | 1 | 15.3 |
| Sulphoraphane | S.10 | 1 | 32.3 |
| Ascorbic acid | C.1 | 1 | 54.1 |
| Epigallocatechin gallate | C.3 | 1 | 57.0 |
| Flavonoid | C.32 | 1 | 41.7 |
| Procyanidin B2 | C.11 | 1 | 55.2 |
| Quercetin | C.15 | 1 | 66.7 |

## IV. Conclusion

Using human SIRT3 and fungal Sir2 as the point of contact, this *in-silico* analysis investigated dietary polyphenols for their capability to modulate aging-oxidative stress pathways within Candida albicans. They are important in the regulation of chromatin remodelling, NAD□ metabolism, and cellular responses to oxidative stress. Molecular docking revealed significant binding affinities of numerous natural substances; quercetin and naringenin emerged as particularly interesting candidates. The potential of quercetin to counteract the reactive oxygen species was evidenced through its major binding contacts with both SIRT3 and Sir2, in addition to its predicted peroxidase inhibition activity following PASS prediction. In addition to expected effects such as membrane integrity agonism, and chlordecone reductase inhibition, naringenin also is significantly binding that suggests a protective role during stressful conditions.

The metabolic importance of these compounds was further supported by SOM based site of metabolism (SOM) predictions; with quercetin and naringenin exhibiting the highest ΔP values at atoms C.15 and 6, respectively for quercetin (Δ=66.2), indicating potential sites of enzyme attack. The good pharmacokinetic properties of both quercetin and naringenin, which includes high gastrointestinal absorption as well as acceptable drug-likeness via the ADME estimations. The polyphenols, quercetin and naringenin in particular could potentially act as modulators of oxidative stress and aging-associated processes in fungal systems based on the integrated computational analyses. This suggests they can be potentially developed as therapeutic or nutraceutical agents.

## Acknowledgement

The authors thank Principal, B.M.S College of Engineering, Management and Department of Biotechnology for providing the necessary facilities to carry out this research work as a part of Major project for final year students.

